# Disease-associated scaffold protein CNK2 modulates PSD size and influences trafficking of new synaptic binding partner TNIK

**DOI:** 10.1101/532374

**Authors:** Hanna L. Zieger, Stella-Amrei Kunde, Nils Rademacher, Bettina Schmerl, and Sarah A. Shoichet

**Affiliations:** Charité-Universitätsmedizin Berlin, Charitéplatz 1, 10117 Berlin, GERMANY

## Abstract

Scaffold proteins are responsible for structural organisation within cells; they form complexes with other proteins to facilitate signalling pathways and catalytic reactions. The scaffold protein connector enhancer of kinase suppressor of Ras 2 (CNK2) is predominantly expressed in neural tissues and was recently implicated in X-linked intellectual disability (ID). We have investigated the role of CNK2 in neurons in order to contribute to our understanding of how CNK2 alterations might cause developmental defects, and we have elucidated a functional role for CNK2 in the molecular processes that govern morphology of the postsynaptic density (PSD). We have also identified novel CNK2 interaction partners and explored their functional interdependency with CNK2. We focussed on the novel interaction partner TRAF2- and NCK-interacting kinase TNIK, which is also associated with ID. Both CNK2 and TNIK are expressed in neuronal dendrites and concentrated in dendritic spines, and staining with synaptic markers indicates a clear postsynaptic localisation. Importantly, our data highlight that CNK2 plays a role in directing TNIK subcellular localisation, and in neurons, CNK2 participates in ensuring that this multifunctional kinase is present at desirable levels at synaptic sites. In summary, our data indicate that CNK2 expression is critical for modulating PSD morphology; moreover, our study highlights a role for CNK2 in directing the localisation of regulatory proteins within the cell. Importantly, we describe a novel link between CNK2 and the regulatory kinase TNIK, and provide evidence supporting the idea that these proteins play complementary roles in the regulation of dendritic spine growth and maintenance.

## Introduction

Scaffold proteins are multi-domain proteins that typically lack enzymatic activity. However, they are crucial in regulating signal transduction cascades: through multiple protein-protein interactions, they organise protein complex formation and ensure spatiotemporal organisation of signalling processes and signal propagation. The scaffold protein connector enhancer of kinase suppressor of Ras (CNK/CNKSR) was first discovered in *Drosophila,* where it was shown to regulate Ras/MAPK signalling by binding to the Ras effector RAF and thereby play an essential role in eye and wing development ^1,2^. Subsequent studies investigating CNKs in various organisms showed that CNK homologues are present across species, ranging *e.g.* from *C. elegans* ^3^ to humans ^1^, and that they likewise influence signalling by acting as scaffolds downstream of Ras ^1^. CNKs possess multiple protein interaction domains, including *e.g.* a sterile alpha motif (SAM), a conserved region in CNK (CRIC), a PSD-95/DLG-1/ZO-1 (PDZ) domain, and a pleckstrin homology (PH) domain, and the domain architecture is essentially conserved throughout the CNK family proteins and also across species.

Since the original discovery of the D-CNK protein in *Drosophila*, it has become clear that CNK family homologues serve as scaffolds for multiple signal cascades: By interaction with various guanine nucleotide exchange factors (GEFs) and GTPase activating proteins (GAPs), they modulate signalling mediated not only by Ras but also via Rho family small GTPases ^4-6^. It has further been shown that CNKs interact with the cytohesin family of ArfGEFs: CNKs facilitate their membrane recruitment, and thereby regulate the insulin signalling pathway ^5,6^. Other studies suggest that CNKs may also regulate cell proliferation and migration by acting as scaffolds directly for the PI3K/Akt ^7^ and JNK ^8^ signalling cascades.

Among the known CNK proteins, mammalian CNK2 isoforms are the proteins that exhibit tissue-specific expression in nervous tissue ^9^. There are studies on how CNK2 isoforms participate in protein-protein interactions ^9-11^, but relatively few studies focus specifically on the scaffold function of CNK2 in neurons and in the brain, despite an increasing number of genetic studies that directly implicate CNK2 mutations in human brain disorders ^12-17^. There is evidence supporting a role for CNK2 in regulating GTPase activity in neurons ^5^, and a recent high-throughput study suggests that CNK2 may be part of the core scaffold machinery that assembles synaptic signalling complexes in the postsynaptic density (PSD) during development ^18^.

Understanding the molecular details of its function during development of nervous tissue will contribute to our knowledge about how CNK2 alterations can cause neurodevelopmental disorders. To explore the function of CNK2, we first examined its specific localisation in neurons and confirmed that it is concentrated at postsynaptic sites ^19^. We next utilised an shRNA-mediated knockdown approach to explore the effects of loss of CNK2 function at these postsynaptic sites, *i.e.* in dendritic spines of glutamatergic neurons. We also took advantage of several CNK2 variants and comparatively investigated their properties in heterologous cells and in dendritic spines. Together, these data suggest a functional role for CNK2 in regulation and maintenance of protein content and morphology of dendritic spines. We also performed a Y2H screen for novel brain-expressed CNK2 interaction partners and identified several protein partners that may participate in the execution of CNK2-mediated regulatory functions in neurons. Subsequent studies focussed on the links between CNK2 and the kinases of the MINK1/TNIK family, for which we discovered a specific interaction with CNK2. We observed a clear functional interdependency between CNK2 and TNIK: subcellular localisation of TNIK is regulated by CNK2 in heterologous cells and in neurons. In summary, our data provide evidence supporting the idea that CNK2 and TNIK/MINK1 family kinases work together in the neuronal environment to ensure proper maintenance and regulation of dendritic spines.

## Results

Mutations in the membrane-associated scaffold protein CNK2 cause developmental disorders; specifically, both deletions ^12-14,17^ and point mutations ^15,16^ that result in loss of function of CNK2 cause X-linked intellectual disability (XLID) that is typically associated with seizures. As CNK2 has its highest expression levels in the brain (^9^; see also www.proteinatlas.org), we were interested particularly in the role of the protein in the development and function of neurons.

### CNK2 is a membrane-associated protein expressed in neurons and enriched at postsynaptic sites

As CNK2 is important for early cognitive development in humans, we first assessed its temporal expression in nervous tissue. Based on western blot of mouse brain lysates we could show that the protein is expressed from postnatal day zero (P0) throughout adulthood **(Fig. 1A)**. We next explored its subcellular localisation. Following ectopic expression of a tagged variant of CNK2 in COS-7 cells, we observed a striking membrane-specific localisation **(Fig. 1B)**. As the commercially available antibodies did not meet expectations, we produced a custom-made antibody against all known CNK2 isoforms. From immunofluorescence experiments on cultured primary rat hippocampal neurons at DIV21 we could see that endogenous CNK2 is enriched at postsynaptic sites in dendritic spines: it co-localises with PSD-95 and shows adjacent staining with the presynaptic marker Synapsin **(Fig. 1C, E, F)**. The postsynaptic localisation of CNK2 was also observed using other methods: following virus-mediated expression of EGFP-tagged CNK2 in hippocampal neurons, we assessed EGFP signal in mature neurons (DIV 21): again we observed enriched signal intensity in spines **(Fig. 1D)**.

**Figure 1:**
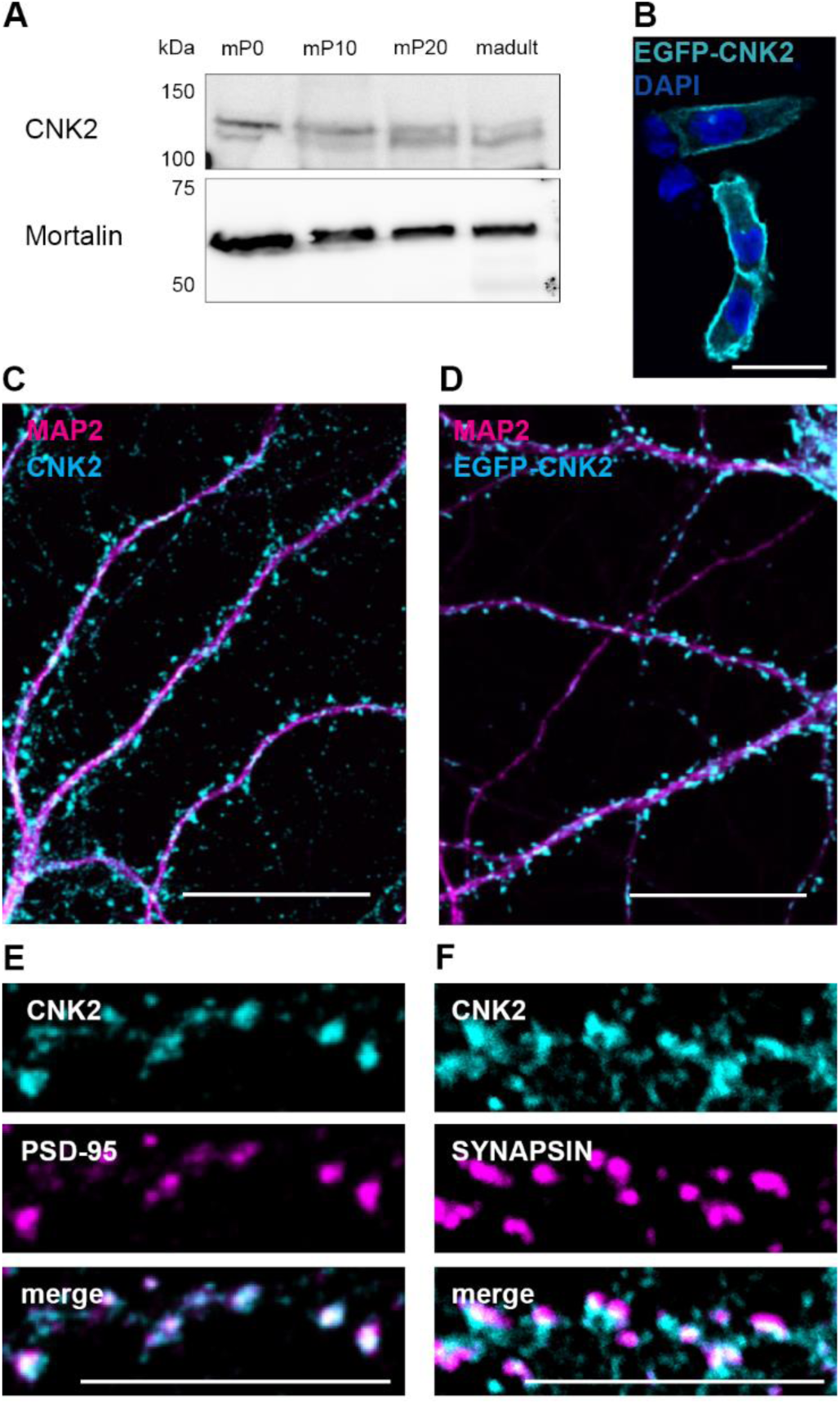
CNK2 is a membrane-associated protein and localised in the postsynapse of neurons. (A) CNK2 is expressed from P0 throughout adulthood in the mouse brain. Whole cell lysates generated from mouse brains at P0, P10, P20 and adult stage were analysed by SDS-PAGE and western blot with anti-CNK2 antibody. Mortalin serves as loading control (expected size 60 kDa). (B) Membrane localisation of overexpressed EGFP-tagged CNK2 (EGFP-CNK2) in COS-7 cells (EGFP-CNK2 cyan; DAPI blue). Scale bar: 20 µm. (C) Localisation of endogenous CNK2 (cyan) and the dendritic marker MAP2 (magenta) in cultured rat hippocampal neurons (DIV21). Endogenous CNK2 is localised in punctate structures along the dendrite. Scale bar: 20 µm (D) Localisation of EGFP-CNK2 (cyan) following lentiviral transduction, and the dendritic marker MAP2 (magenta) in cultured rat hippocampal neurons (DIV21) Scale bar 20 µm. (E) CNK2 (cyan) colocalises with the postsynaptic marker PSD-95 (magenta). (F) CNK2 (cyan) shows adjacent staining with the presynaptic marker Synapsin (magenta). Scale bar: 10 µm.

### Loss of CNK2 influences the size of the postsynaptic density

Given that complete loss of CNK2 was implicated in brain disorders, together with our observation that CNK2 is expressed in neurons, in particular in dendrites and at postsynaptic sites, we were interested to explore the idea that dendritic spines might be affected by loss of CNK2. We took advantage of lentivirus-mediated gene delivery of an shRNA ^5^ to specifically knockdown endogenous CNK2 in primary neurons (**Supplemental Fig. 1**). Eighteen to nineteen days after transduction, neurons were fixed and stained for the postsynaptic marker Homer and the dendritic marker MAP2 **(Fig. 2A)**. As a reflection of PSD size, we quantified the endogenous Homer content in mature spines (DIV 21)^20,21^. Blinded analysis of the Homer immunofluorescence signal area using the “Analyze Particles” tool (FIJI/ ImageJ) ^22^ enabled a quantitative comparative analysis of Homer immunofluorescence signal intensity and area in neurons expressing either CNK2 shRNA or control shRNA **(Fig. 2B)**. This analysis revealed a clear reduction of PSD size in CNK2 knockdown neurons (overall reduction of 15%; p < 0.0001) **(Fig. 2C)**. We also performed a comparative analysis of the spine density in neurons infected with either CNK2 knockdown or control shRNA: here we did not observe substantial differences (data not shown).

**Figure 2:**
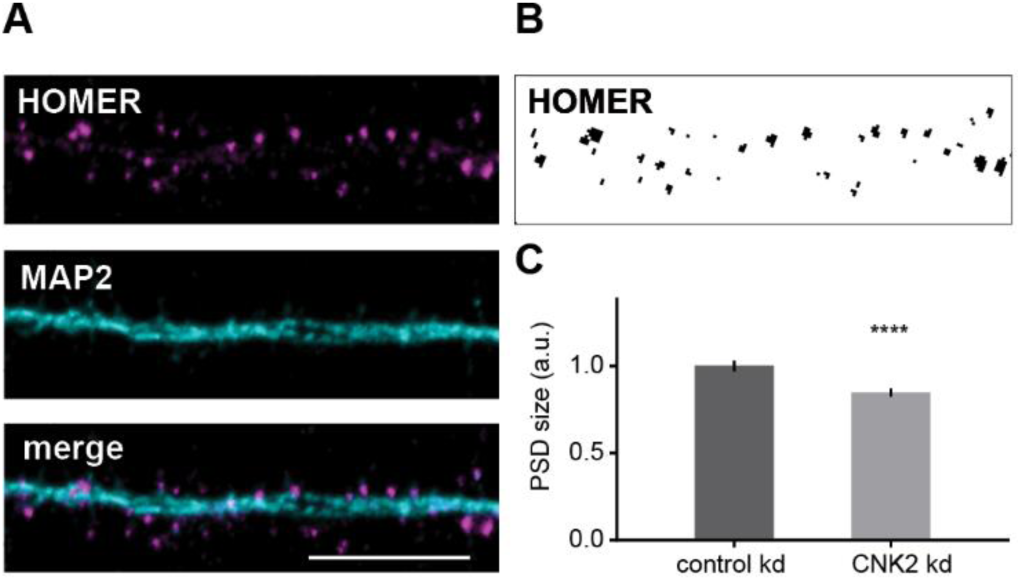
Loss of CNK2 reduces PSD size. (A) Cultured rat hippocampal neuron infected with lentivirus transducing shRNA stained for the postsynaptic marker Homer (magenta) and dendritic marker MAP2 (cyan). Scale bar: 10 µm. (B) Sample image of endogenous Homer thresholded for analysis of Homer particle size. (C) PSD size, represented by the signal of the postsynaptic marker protein Homer (arbitrary units), was analysed using the “Analyze Particles” tool (FIJI/ Image). Data were normalised to the mean of the control. Graph represents the mean ± SEM (n= 757-1100 spines from 7-10 cells, N=3 cultures): control (scrambled shRNA) = 1 ± 0.03, CNK2 knockdown = 0.85 ± 0.02; Mann-Whitney test, p < 0.0001.

### Expression of a CNK2 variant that does not bind to the membrane affects PSD size

We observed that loss of CNK2 influences PSD size (see **Fig. 2**). We also validated that wild-type CNK2 is membrane-associated in heterologous cells (**Fig. 1B**). In order to explore the functional importance of its membrane localisation, we generated an EGFP-tagged CNK2 deletion construct that lacks the C-terminal region including the PH domain (CNK2ΔPH; see **Supplemental Fig. 2** and also **Fig. 5A** for overview of CNK2 domain architecture). Following ectopic expression of this mutant, we observed a clear loss of membrane association (**Supplemental Fig. 2**), which is in line with biochemical studies indicating that PH domains typically bind to phosphatidylinositol lipids in biological membranes ^23^. We expressed this construct in primary hippocampal neurons and compared its expression with EGFP-tagged wild-type CNK2 regarding its influence on PSD size. As done for our CNK2 knockdown neurons, we utilised a quantitative comparative immunofluorescence approach to analyse protein content in dendrites and spines (**Fig. 2**). We first assessed the general expression of wild-type EGFP-CNK2 and EGFP-CNK2ΔPH in dendrites by comparing EGFP signal intensity relative to the dendritic marker MAP2 (**see Fig. 3A**), and we observed dramatic differences (for CNK2ΔPH, normalised dendritic signal intensity was approximately one-third that of the wild-type; see **Fig. 3B)**. To make sure that this was not an effect of globally reduced expression of the truncated CNK2 variant, we analysed whole cell lysates from neurons overexpressing the CNK2 variants by western blot and observed comparable expression levels for EGFP-CNK2 and EGFP-CNK2ΔPH **(Supplement Fig. 3B)**. These observations suggest that this difference do not simply reflect reduced total expression of EGFP-CNK2ΔPH but rather specifically reflect an altered localisation.

**Figure 3:**
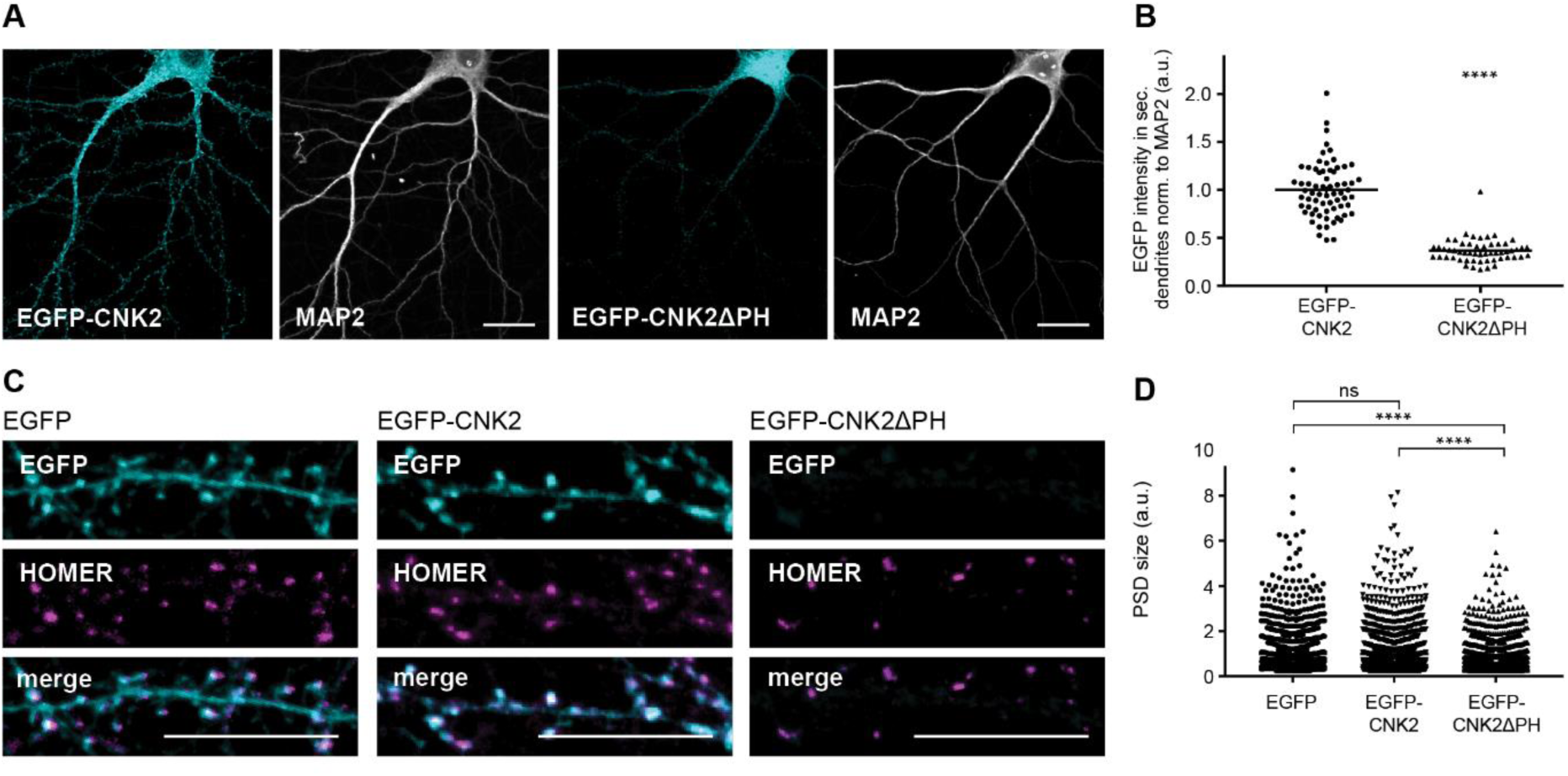
Expression of a CNK2 variant that does not bind to the membrane affects PSD morphology. (A) Representative images of primary neurons expressing either EGFP-CNK2 (cyan) or EGFP-CNK2ΔPH (cyan) co-stained for endogenous MAP2 (grey). Scale bar: 20 µm. (B) Quantification of amount of EGFP-CNK2 and EGFP-CNK2ΔPH based on EGFP-signal intensity normalised to MAP2 signal intensity in regions of interest (ROIs) along secondary dendrites. Data were normalised to the mean of the control. Graph represents mean ± SEM (arbitrary units, a.u.) of EGFP intensity (n= 55-67 ROIs in 11-14 neurons, N=3 cultures): EGFP-CNK2 = 1 ± 0.04, EGFP-CNK2ΔPH = 0.37 ± 0.02, data were analysed by Mann-Whitney test, p < 0.0001. (C) Neurons expressing EGFP, EGFP-CNK2 or EGFP-CNK2ΔPH stained for EGFP (cyan) and Homer (magenta). Scale bar: 10 µm. (D) Postsynaptic density (PSD) size upon expression of EGFP, EGFP-CNK2 or EGFP-CNK2ΔPH. PSD size, represented by the signal of the postsynaptic marker protein Homer (a.u.), was analysed using the “Analyze Particles” tool (FIJI/ImageJ). Data were normalised to the mean of the control. Graph represents the mean ± SEM (n= 1555-2008 spines in 14-18 neurons, N=4 cultures); EGFP = 1 ± 0.02, EGFP-CNK2 = 0.99 ± 0.02 or EGFP-CNK2ΔPH = 0.81 ± 0.02; data were analysed by Kruskal-Wallis test, followed by Dunn’s multiple comparison test: EGFP vs. EGFP-CNK2: p = 0.343, EGFP vs. EGFP-CNK2ΔPH: p < 0.0001; EGFP-CNK2 vs. EGFP-CNK2ΔPH: p < 0.0001.

**Figure 4:**
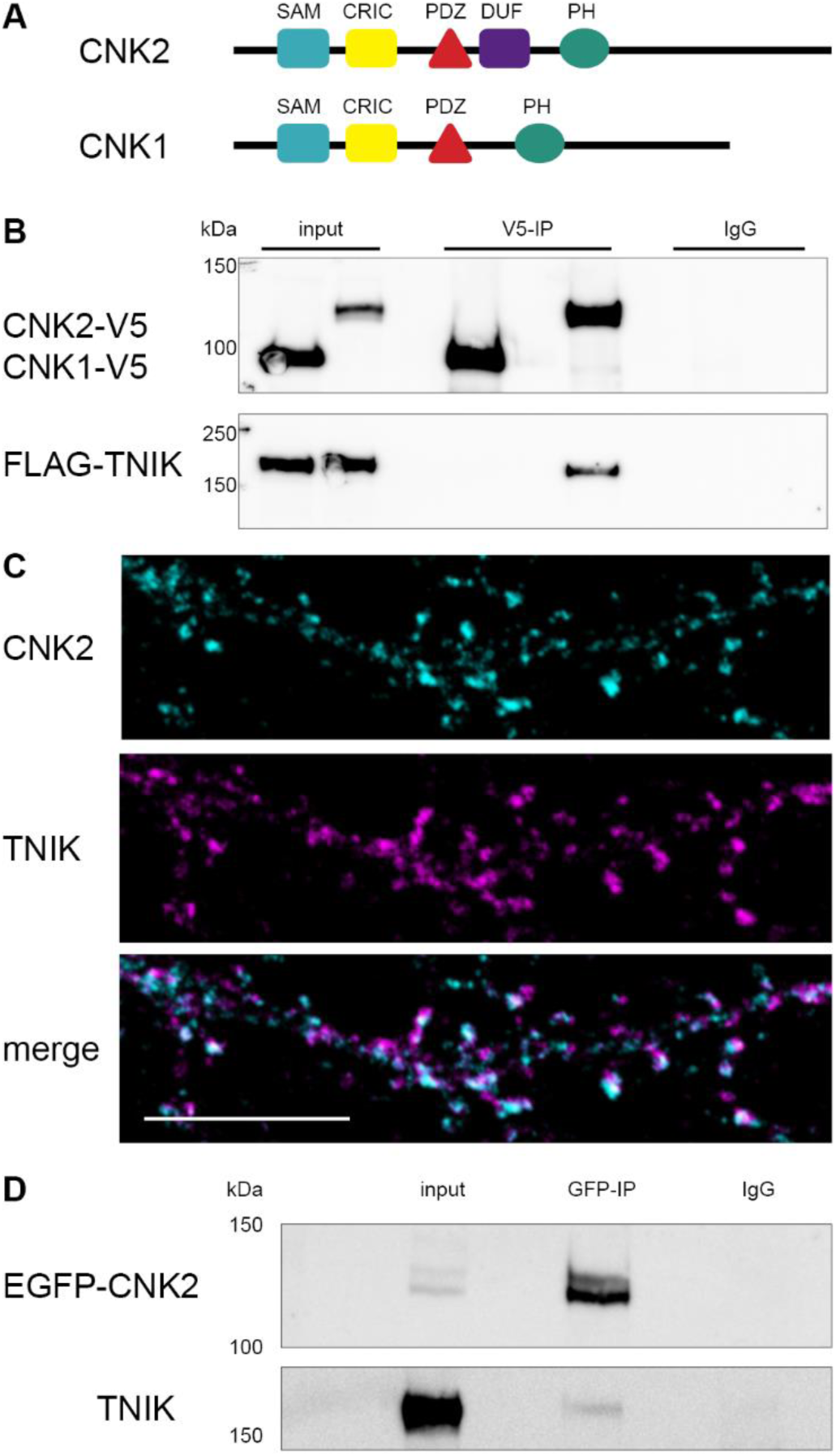
CNK2 interacts with TNIK and the two proteins co-localise in dendritic spines. (A) Scheme of CNK1 and CNK2 domain architecture (SAM, sterile alpha motif; CRIC, conserved region in CNK; PDZ, PSD-95/DLG-1/ ZO-1); DUF1170, domain of unknown function; PH, pleckstrin homology). (B) CNK2 specifically interacts with TNIK. Coimmunoprecipitation experiment with CNK1-V5, CNK2-V5 and FLAG-TNIK expressed in COS-7 cells. Proteins were immunoprecipitated with either anti-V5 (mouse) antibody or mouse IgGs as a negative control. Proteins were detected by western blot with anti-V5 (CNK2) and anti-FLAG (TNIK) antibodies. (C) Representative image of a primary neuron (DIV 21) showing endogenous CNK2 (cyan) and endogenous TNIK (magenta) in dendritic spines. Scale bar: 10 µm (D) Co-immunoprecipitation experiment of EGFP-CNK2 with endogenous TNIK expressed in neurons. Protein was immunoprecipitated with either anti-GFP (mouse) or mouse IgGs as a negative control. Proteins were detected by western blot with anti-GFP (CNK2) and anti-TNIK antibodies. Input control (lysate) is shown on the left.

**Figure 5:**
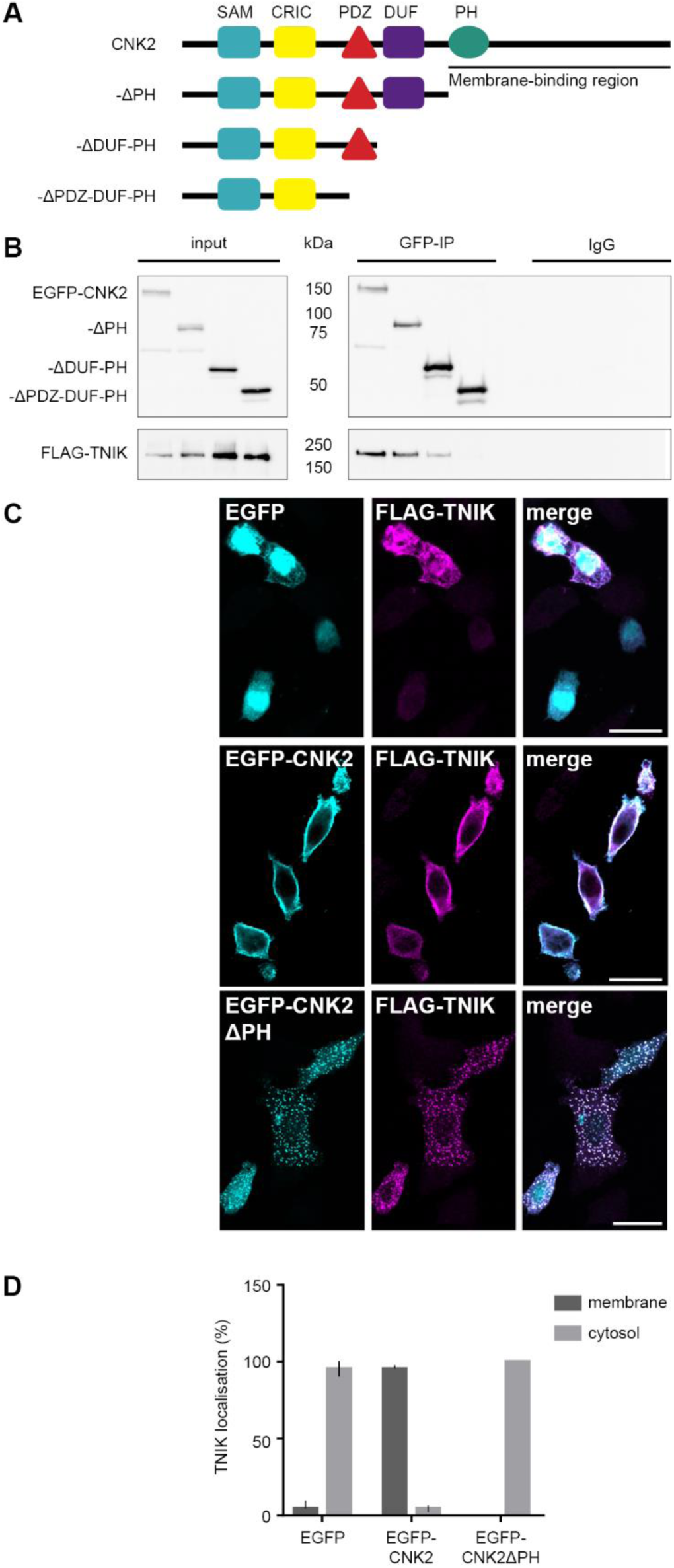
TNIK localisation is regulated by CNK2. (A) Scheme of CNK2 truncation variants (for detailed description of domains see Fig. 4A legend). (B) Co-immunoprecipitation experiments of EGFP-CNK2 variants overexpressed in COS-7 cells together with FLAG-TNIK. Proteins were immunoprecipitated with either anti-GFP (mouse) antibody or mouse IgGs as a negative control. Pull-down control and coprecipitated proteins were analysed by western blot with anti-GFP (IP) and anti-FLAG (coIP) antibodies. Input control (lysate) is shown on the left. (C) TNIK localisation is influenced by CNK2 in heterologous cells. Representative immunofluorescence experiments in COS-7 cells overexpressing FLAG-TNIK together with EGFP as control (upper panel), EGFP-CNK2 (middle panel) or EGFP-CNK2ΔPH (lower panel). Left panels show EGFP only or EGFP-tagged CNK2 variants (cyan), middle panels shows FLAG-TNIK (magenta), and the right lane shows merged channels. Scale bar: 20 µm. (D) For quantification of TNIK localisation in COS-7 cells reflecting the experiments shown in (C), images expressing both proteins, were randomised and classified according to cytosolic or membranous TNIK localisation. Co-expression of CNK2 recruits the main fraction of FLAG-TNIK to the membrane. Upon co-expression of EGFP-tagged CNK2 variants lacking the membrane binding region, FLAG-TNIK shows essentially no membranous localisation. Data used for quantification include data for a total of 73-114 cells per condition, imaged from three independent experiments.

Next, we comparatively assessed expression of the postsynaptic protein Homer **(Fig. 3C)**. Again, following blinded analysis of Homer in spines, we observed a significant reduction of Homer content in the dendritic spines of neurons expressing EGFP-CNK2ΔPH when compared to neurons expressing the wild-type EGFP-CNK2 or EGFP alone **(Fig. 3D)**. In summary, we conclude that, compared to neurons expressing EGFP-CNK2, neurons expressing the non-membrane-associated EGFP-CNK2ΔPH have a reduced PSD size. This result indicates that – as we observe for CNK2 loss of function (**Fig. 2**) – mis-localisation of CNK2 also interferes with regulation of the size of the PSD.

### CNK2 interacts with TNIK and the two proteins co-localise in dendritic spines

To further investigate the role of CNK2 in neurons, we next focussed on novel CNK2 interaction partners, which we identified via a yeast-two-hybrid (Y2H) screen of cDNAs from adult mouse brain. The list of interactors included proteins with diverse functions, and a significant fraction of positive hits were signalling proteins **(see Supplemental Table 1)**. Importantly, our list included the Rho-GTPase activating protein Vilse/ARHGAP39, which has previously been shown to interact with CNK2 ^5^, and thus served as a positive control for our Y2H screen. Two additional proteins identified in our screen, the kinases TNIK and MINK1, were of special interest to us, in part because they have both been implicated in the regulation of neuron structure and glutamate receptor function ^24^. In addition, TNIK was recently shown to be associated with intellectual disability in patients ^25^, and TNIK knockout mice exhibit cognitive impairment and hyperactivity ^26^. Moreover, this protein was recently investigated for its role in the regulation of protein complexes in PSDs ^18^.

MINK1 and TNIK have common domain architecture but exhibit minimal sequence homology outside of their conserved domains. We demonstrated that CNK2 interacts with both TNIK and MINK1 in co-immunoprecipitation assays **(Fig. 4, Supplement Fig. 4).** Interestingly, the ubiquitously expressed CNK1 protein, which shares several interaction domains with CNK2 **(Fig. 4A)**, did not interact with TNIK or MINK1 in comparable assays **(Fig 4B, Supplement Fig. 4A)**, suggesting that the CNK2-TNIK and CNK2-MINK1 interactions may have CNK2-specific functions that are important in neurons.

Subsequent cell-based studies focussed on the CNK2-TNIK interaction. We first investigated whether CNK2 and TNIK indeed reside in the same subcellular compartments. Using antibodies to the endogenous proteins, we could show by immunofluorescence on cultured primary rat hippocampal neurons (DIV21) that CNK2 and TNIK co-localise in dendritic spines **(Fig. 4C)**, providing further support for the idea that these two proteins can indeed function together at postsynaptic sites. We also confirmed that endogenous TNIK binds CNK2 in neural tissue: following immunoprecipitation of EGFP-CNK2 expressed in primary rat hippocampal neurons at DIV21, we could detect coprecipitated TNIK by western blot analysis **(Fig. 4D)**.

### TNIK localisation is regulated by CNK2

In a subsequent set of experiments, we narrowed down the region within CNK2 relevant for TNIK or MINK1 binding. Using a set of EGFP-tagged CNK2 deletion constructs with C-terminal truncations of various lengths **(Fig. 5A)**, we demonstrated via coimmunoprecipitation that loss of the C-terminal region harbouring the PH domain had no effect on TNIK binding. Deletion of both PH and DUF domains, however, did affect binding affinity. Most notably, when the PDZ domain was deleted together with the PH and DUF domains, binding to TNIK and MINK1 was completely abolished **(Fig. 5B, Supplement Fig. 4B)**. Together, these results indicate that the region including the PDZ and the DUF domain is critical for CNK2 binding to TNIK and MINK1, and that a CNK2 variant lacking only the PH domain / C-terminus responsible for membrane localisation is still capable of efficient binding to these new interacting proteins.

We next took advantage of this MINK/TNIK-binding deletion construct (EGFP-CNK2ΔPH) to explore the idea that CNK2 might functionally influence TNIK. We observed earlier that full-length CNK2 exhibits a clear membrane localisation in heterologous cells **(Fig. 1B)**. Immunofluorescence of ectopically expressed FLAG-TNIK together with EGFP as a control in COS-7 cells indicated that TNIK is not enriched at the membrane **(Fig. 5C, upper panel)**. However, upon co-expression with full-length CNK2, TNIK was observed at the membrane in most cells **(Fig. 5C, middle panel)**, suggesting that CNK2 might play an important role in regulating the subcellular localisation of TNIK. In order to confirm the specificity of this observation we took advantage of the CNK2ΔPH mutant described earlier **(see Fig. 5A)**. Upon co-transfection of EGFP-CNK2ΔPH, TNIK was no longer at the membrane **(Fig. 5C, lower panel)**, suggesting that binding to wild-type CNK2 is indeed a mechanism that facilitates TNIK membrane localisation. A comparable experiment with CNK2 and MINK1 indicates that the same is true for this protein-protein interaction **(Supplement Fig. 4C)**, *i.e.* our data suggest that CNK2 is capable of influencing the membrane localisation of both kinases.

We next quantified these results following blinded analysis of TNIK membrane localisation in cells co-transfected with TNIK and either wild-type or mutant CNK2 proteins **(Fig. 5D)**. This quantification clearly demonstrates that TNIK localisation is influenced by CNK2 localisation in heterologous cells.

### Mis-localisation of CNK2 also influences TNIK localisation in neurons

To gain further insight into the CNK2-mediated regulation of TNIK in neurons, we examined the precise neuronal localisation of TNIK in neurons expressing either wild-type EGFP-CNK2 or EGFP-CNK2ΔPH. Immunofluorescence of EGFP-tagged proteins in neurons confirmed that – as in heterologous cells – EGFP-CNK2ΔPH exhibited an aberrant localisation in neurons: compared to wild-type CNK2, this truncated version was expressed at reduced levels in dendrites; instead, it accumulated in the soma and in the nucleus, which was not the case for full-length EGFP-CNK2 **(Supplement Fig. 3A)**. As we showed earlier **(Fig. 1)**, full-length CNK2 is observed primarily in dendrites and in spines. We next assessed TNIK localisation in neurons expressing either full-length or truncated CNK2 variants. Cultured rat hippocampal neurons expressing either EGFP alone, EGFP-CNK2 or EGFP-CNK2ΔPH were analysed with regard to TNIK content and localisation (**Fig. 6A**). We normalised endogenous TNIK signal to the signal for the dendritic marker MAP2 and used a quantitative immunofluorescence approach that involved a blinded selection and analysis of the images. On average, the intensity of endogenous TNIK in secondary dendrites was reduced significantly by 18% (p=0.0001) or 14% (p=0.0027) in neurons expressing EGFP-CNK2ΔPH compared to neurons expressing EGFP or EGFP-CNK2, respectively **(Fig. 6B)**. These data suggest that mis-localisation of CNK2ΔPH, which binds to TNIK, indeed seems to trap TNIK in the soma, thereby resulting in reduced total TNIK in the dendrites. In our studies, in line with other publications ^27^, we show that endogenous TNIK is naturally localised in the postsynapse of wild-type neurons. We explored the idea that loss of CNK2 might influence TNIK localisation but we could not observe a significant difference of TNIK content in dendrites following virus-mediated shRNA knockdown of CNK2 in neurons (data not shown). However, the amount of endogenous TNIK in the dendrites is clearly reduced in neurons expressing the mis-localised truncated CNK2 (CNK2ΔPH) **(Fig. 6)**. This result is in line with our data from heterologous cells, and suggests that this mutant is capable of interfering with normal CNK2-mediated regulation of TNIK localisation in neurons.

**Figure 6:**
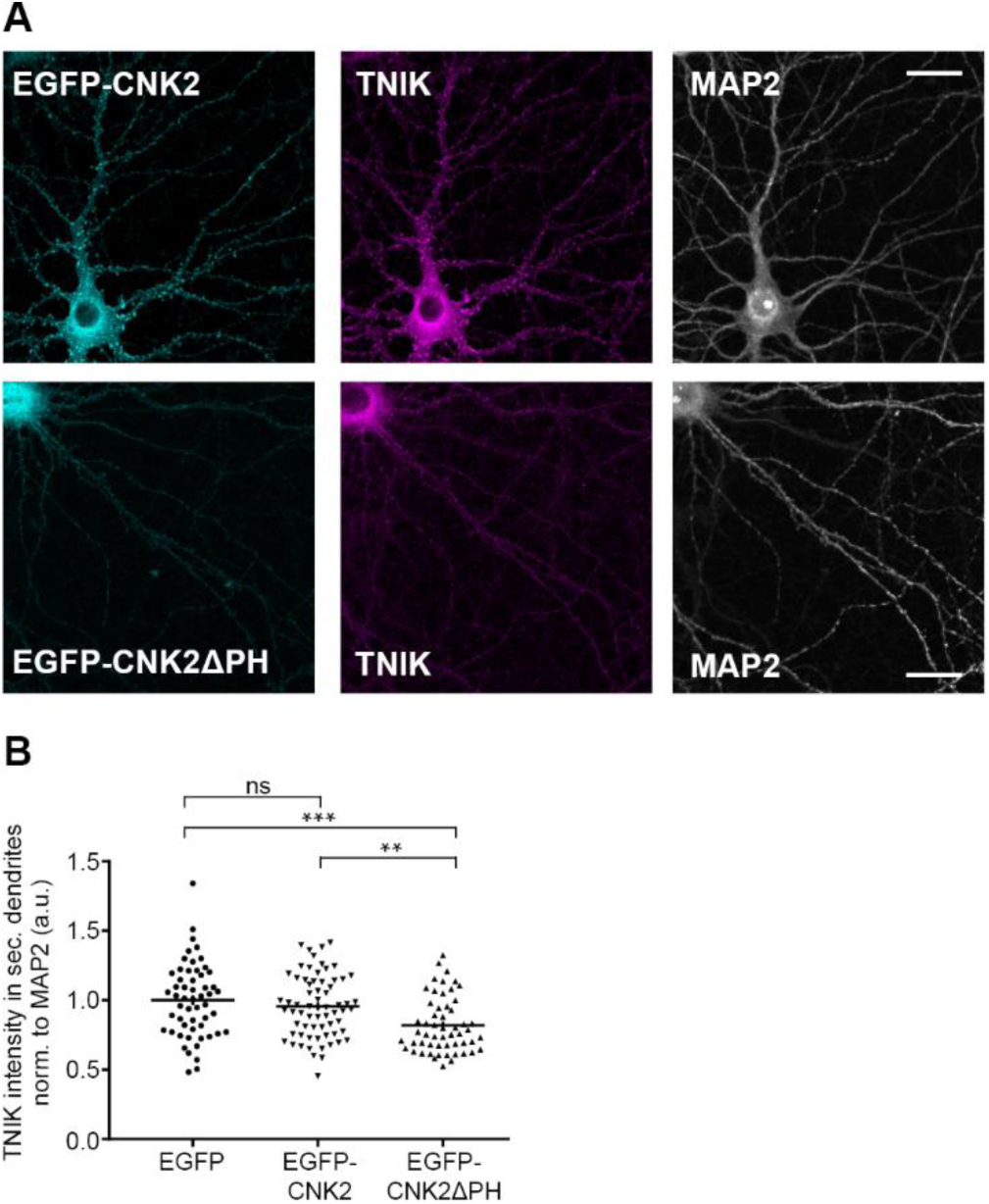
Mis-localisation of CNK2 also influences TNIK localisation in neurons. (A) Images of primary neurons expressing EGFP-CNK2 (cyan) or EGFP-CNK2ΔPH (cyan) co-stained for endogenous TNIK (magenta) and MAP2 (grey). Scale bar: 20 µm (B) Co-expression of EGFP-CNK2ΔPH causes a reduction of TNIK in secondary dendrites compared to EGFP or EGFP-CNK2. TNIK intensity was measured in ROIs along secondary dendrites and normalised to MAP2 (a.u.). Data were normalised to the mean of the control. Graph represents mean ± SEM of TNIK intensity (n= 55-67 ROIs in 11-14 neurons, N=3 cultures): EGFP = 1 ± 0.036, EGFP-CNK2 = 0.96 ± 0.027, EGFP-CNK2ΔPH = 0.82 ± 0.027, data were analysed by ordinary one-way ANOVA, followed by Dunnett’s post-hoc test: EGFP vs. EGFP-CNK2: p = 0.474; EGFP vs. EGFP-CNK2ΔPH: p = 0.0001; EGFP-CNK2 vs. EGFP-CNK2ΔPH: p = 0.0027

## Discussion

Patients with CNK2 mutations exhibit an array of neurocognitive symptoms, ranging from mild intellectual disability and language delay to more severe and general delayed cognitive and motor development. Seizures are also present in most patients ^15,16^. Clearly, loss of functional CNK2 has detrimental consequences for normal development of functional neural networks; however, the precise molecular mechanisms by which CNK2 mutations cause disease have not been elucidated to date. Our results validate that CNK2 is expressed in neurons and enriched at postsynaptic sites. Further studies revealed a set of novel binding partners for CNK2. Interestingly, the proteins identified were not structural proteins that have been previously shown to participate in regulating PSD size. Instead, our list of interacting proteins consisted predominantly of regulatory proteins, including, for example the Rho-GTPase activating protein Vilse/ARHGAP39, which has previously been investigated for its role in mediating CNK2 function ^5^, and the regulatory kinases MINK1 and TNIK. We focussed on the new CNK2-interacting kinase TNIK which, like CNK2, was recently implicated in cognitive disorders ^25^. TNIK is a well-characterised signalling molecule that plays a decisive role in the activation of multiple signal cascades ^28,29^. More recently, it has been shown that it is concentrated in dendritic spines ^27^, and that it plays a role, together with MINK1, in the regulation of AMPAR trafficking ^24^. We confirmed that CNK2 and TNIK exhibit coexpression in dendrites and at postsynaptic sites, and taking advantage of TNIK-binding CNK2 variants that exhibit aberrant subcellular localisation, we demonstrated that CNK2 directly modulates neuronal TNIK, and thus provide strong support for the idea that TNIK and CNK2 participate in common pathways that may be critical for the observed CNK2-mediated regulation of PSD size.

Interestingly, it has been proposed by others that TNIK is involved in the regulation of synaptic protein degradation ^30^. Also relevant is that both p120-catenin and delta-catenin have recently been identified as direct TNIK substrates ^29^. Both of these catenins interact with cadherins at synapses, and multiple studies demonstrate that their regulation is an important factor in modulating dendritic spine architecture ^31-34^. Moreover, TNIK is an important regulator of JNK signaling ^28,35,36^, which may also have implications for regulating synaptic protein content ^37,38^. It is thus plausible that alterations in synaptic TNIK expression levels or activity might contribute indirectly to changes in the expression of multiple other synaptic proteins and thereby regulate dendritic spine architecture.

Given that studies in humans and mice highlight a role for CNK2 and TNIK in related disorders, combined with the fact that diverse studies demonstrate that TNIK plays an established role in neuronal differentiation and network formation, we propose that this novel CNK2 binding partner might work in concert with CNK2 during normal neural development. Although we could not measure significant changes in TNIK synaptic expressions levels upon knockdown of CNK2, our studies indeed highlight a functional interdependence of these two proteins with regard to membrane trafficking and thereby illuminate a putative regulatory cascade that may underlie the defects responsible for the observed cognitive dysfunction in patients with CNK2 alterations. Given their physical interaction, it is also plausible that TNIK regulatory activity may be influenced by CNK2. Finally, it is important to note that while we focus here on the CNK2-interacting proteins of the TNIK/MINK family, it is likely that several other regulatory proteins are also affected by disease-associated changes in CNK2 expression. Future studies aim to explore in greater depth how diverse CNK2-mediated signal cascades participate in the cellular processes that govern neurotransmission and proper network formation.

## Materials and Methods

### Constructs

Cnk2 (Accession number: NM_177751) was amplified from cDNA generated from mouse brain. The pEGFP-C1-Cnk2 construct (accession number: XM_011247826.1) was cloned using the primers Cnk2-SalI-fw (aagtcgacatggctctgataatggaaccgg) and Cnk2-BamHI-rv (ttggatccttagcttttctctccaac). For the truncated variants, the forward primer Cnk2-SalI-fw was used together with Cnk2ΔPH-rv (ttggatccttagccaagatcctttca) for pEGFP-C1-CNK2ΔPH, Cnk2ΔDUF-PH-rv (ttggatccttaaagcatgctctgagg) for pEGFP-C1-CNK2ΔDUF-PH and Cnk2ΔPDZ-DUF-PH-rv (ttggatccttattccaggtgagcaga) for pEGFP-C1-CNK2 ΔPDZ-DUF-PH. PCR products were cloned into pEGFP-C1 (Clontech) using BamHI and SalI. EGFP-CNK2 and truncated variants were cloned into a lentiviral shuttle vector under the control of a human synapsin-1 promoter using BamHI and SalI restriction sites. Mouse Cnk2 was subcloned into pBudCE4.1 using pBud-Cnk2-Notl-fw (aagcggccgcatggctctgataatggaa) and pBud-Cnk2-KpnI-rv (ttggtaccgcttttctctccaacgtt). Rat Cnk1 (accession number: BC099788) was amplified from cDNA-clone (Clone ID: 7934518, Source Bioscience) and subcloned into pBudCE4.1 using the primers rCnk1NotI-fw (aagcggccgcatggagcccgtggag) and rCnk1-BglII-rv (aaagatctgggaggtcaggaggtt).

Vectors expressing FLAG-TNIK and FLAG-MINK1 were generous gifts from Natasha Hussain, (Hussain et al, 2011). FLAG MINK1-C (AA 534-1310) was subcloned using the primers Mink1-C-XbaI-fw (aatctagaatgcagcagaactctccc) and Mink1-rv (gtaaccattataagctgc).

### Co-immunoprecipitation, SDS-PAGE and western blot

COS-7 cells were maintained in DMEM (Lonza) supplemented with 10 % FBS (Sigma), 2 mM L-glutamine and penicillin/streptomycin at 37 °C with 5 % CO2. Transient transfections were done using Lipofectamine 2000 (Invitrogen) according to the manufacturer’s recommendations. Transfected COS-7 cells (T75 flask) were harvested 18-20 hours post transfection in PBS using a cell scraper and pelleted by centrifugation at 1200 × *g*. COS-7 cell pellets were lysed in 1 ml lysis buffer (50 mM Tris pH 7.5, 100 mM NaCl, 1% Triton X) and lysed using a 30G needle. Lysates (1 ml) were cleared by two centrifugations for 10 min at 20817 × *g* and incubated with the appropriate antibody for 3 hours at 4°C. Lysates were cleared by centrifugation for 10 min at 20817 × *g*. Supernatants were incubated with 30 µl Protein G-Agarose (Roche) per ml lysate for 1 hour at 4 °C and washed three times with lysis buffer. Immunocomplexes were collected by centrifugation, denatured, and analysed by SDS-PAGE and western blot (semi-dry blotting system, Bio-Rad). PVDF-membranes (Bio-Rad) were blocked (PBS, 0.1 % Tween 20, 5 % dry milk) and incubated overnight with primary antibody. Membranes were incubated for 1 hour at 4 °C with the respective horseradish peroxidase (HRP)-conjugated secondary antibody. If the primary antibody used was conjugated to HRP no secondary antibody was added. Western Lightning Plus-ECL was used to visualise the signal on the blot and recorded with Image Quant (LAS4000Mini, GE Healthcare). To detect other proteins of interest on the same membrane, the membrane was incubated overnight at 4°C in blocking buffer containing 0.1 % sodium azide with subsequent primary and secondary antibody as described before. For coIP from cultured neurons, infected neurons were washed with warm PBS, subsequently lysed in lysis buffer and further treated as described above.

### Neuron culture

For primary rat hippocampal neuronal cultures, embryonic E18 Wistar rats were used. Following decapitation, hippocampi from embryos were isolated and collected in ice-cold DMEM (Lonza). Neurons were separated using Trypsin/EDTA (Lonza) at 37 °C for 5 min. After stopping the reaction with 10 % FBS (Biochrom) in DMEM and subsequent washing in DMEM to remove trypsin, the hippocampal tissue was suspended in neuron culture medium (Neurobasal A supplemented with B27 and 0.5 mM glutamine) and further dissociated mechanically. For immunofluorescence, neurons were plated onto glass coverslips (d=18 mm) coated with a mixture of poly-D-lysine (Sigma) and Laminin (Sigma) in PBS at a density of 1.5×10^5^ cells per 12-well. For lysates, plates were coated as described before and cells were plated at a density of 7.5 x10^5^ cells per 6 well. Cell debris was removed after healthy neurons adhered (50 minutes post-plating), and neurons were maintained at 37 °C with 5% CO_2_ in neuron culture medium.

### Lenti-viral infection and Immunofluorescence

Cultured neurons on glass coverslips were infected at DIV3 with lentivirus transducing CNK2 shRNA/ control shRNA and at DIV10 for expression of EGFP, EGFP-CNK2 or EGFP-CNK2ΔPH. At DIV23-24, neurons were fixed in 4 % PFA in PBS for 10 min. After washing with PBS the cells were permeabilised in 0.2 % Triton-X in PBS for 5 min, washed again with PBS and blocked with 4 % bovine serum albumin (BSA) in PBS for 1 h at room temperature. Cells were incubated with primary antibodies in 4% BSA in PBS at 4 °C overnight, washed with PBS and subsequently incubated with secondary antibodies in blocking solution. After washes in PBS, coverslips were dipped in deionized water and mounted with Fluoromount-G (Southern Biotech).

### Imaging

Samples were randomised before analysis. Images were acquired with a Leica laser-scanning confocal microscope (Leica TCS-SP5 II) using the 63x immersion oil objective. Total z-stack range of 2 µm was set with a 0.4 µm inter-stack interval and used in a maximal z-stack projection for further analysis.

### Analysis

Analysis was done randomised using FIJI/ ImageJ software (Version 1.52g) ^22^. For determination of distribution of EGFP-CNK2 and EGFP-CNK2ΔPH (cyan) and its effect on TNIK (magenta), regions of interest (ROI) were defined along secondary dendrites, 4-6 ROIs per neuron. Fluorescence intensity per ROI was measured for all channels. Intensity for EGFP and TNIK signal was normalised to intensity of MAP2 signal (grey, 405) per ROI. Measured EGFP signal intensity of EGFP-CNK2 and EGFP-CNK2ΔPH was normalised to the mean of EGFP-CNK2 signal. For TNIK distribution, every value was normalised to the mean of TNIK in the control situation (EGFP only). For spine analysis, dendrites were imaged as described before. PSD size, represented by the signal of the postsynaptic marker protein Homer was analysed using the “Analyze Particles” tool of FIJI. For each experiment, the threshold for the Homer staining representing the PSD size was set according to experimenters’ discretion. Every spine measured was normalised to the median Homer area of the control condition (EGFP).

### Crude synaptosome preparation

For the crude synaptosome preparation, cultured neurons were treated with Syn-PER Synaptic Protein Extraction Reagent (Thermo Scientific) according to the manufacturer’s recommendations and analysed by SDS-PAGE and western blot.

### Antibodies

Primary antibodies: anti-CNK2 (guinea pig, Eurogentec, custom-made), anti-CNK2 (rabbit, Sigma/Atlas, HPA 001502), anti-HOMER-1/2 (guinea pig, Synaptic Systems, 160004), anti-MAP2 (guinea pig, Synaptic Systems, 188004), anti-MAP2 (mouse, Millipore, 05-346), anti-Mortalin (mouse, Antibodies Inc., 75-127), anti-PSD-95 (mouse, Antibodies Inc., 75-028), anti-SYNAPSIN-1 (rabbit, Synaptic systems, 106 103), anti-TNIK (mouse, Santa Cruz, sc-136103), anti-FLAG-HRP (mouse, Sigma), anti-GFP (chicken, Abcam, ab13970), anti-GFP (mouse, Roche, 11814460001), anti-GFP (goat, Abcam, ab6673), normal mouse IgG (sc-2025, Santa Cruz), anti-V5 (mouse, Invitrogen, R960-25), anti-V5 (rabbit, Millipore, AB3792).

Secondary antibodies: anti-mouse-HRP (Dianova, 115-035-003), anti-rabbit-HRP (Dianova, 111-035-003), anti-goat-HRP (Santa Cruz, sc-2020), anti-guinea pig Alexa Fluor 405 (Abcam, ab175678), anti-mouse Alexa Fluor 405 (Invitrogen, A-31553), anti-guinea pig Alexa Fluor 488 (ThermoFisher, A-11073), anti-chicken Alexa Fluor 488 (Jackson Immuno Research, 703-545-155), anti-rabbit Alexa Fluor 568 (Life Technologies, A-11036), anti-mouse Alexa Fluor 568 (Life Technologies, A-11031).

### Custom-made CNK2 antibody

The custom-made CNK2 antibody used in this study was produced by Eurogentec. It was raised in guinea pig against a KHL-conjugated peptide representing CNK2 amino-acids 727-741 and affinity matrix purified. The peptide is present an all known CNK2 isoforms.

### Yeast-Two-Hybrid

The Y2H screen (Hybrigenics ULTImate Y2H) was performed by HYBRIGENICS (Paris, France), using full-length CNKSR2 (mus musculus, Gene ID: 245684, aa 1-1032), cloned into pB27 (N-LexA-bait-C fusion) with an adult mouse brain cDNA library.

### Ethical approval

All animals used in this study were treated according to the German regulations and approved by the ‘Landesamt für Gesundheit und Soziales’ (LaGeSo; Regional Office for Health and Social Affairs) in Berlin on the use of animals for research purposes and sacrificed under the permit T0280/10.

## Acknowledgements

We are very grateful for technical assistance from Melanie Fuchs. We also thank the NeuroCure Virus Core Facility (VCF) at the Charité-Universitätsmedizin Berlin for providing viruses and laboratory space for infection. This work was supported by the ‘Deutsche Forschungsgemeinschaft’ (DFG) Project SH 650/2, the DFG NeuroCure Cluster of Excellence (EXC257), and DFG Collaborative Research Centers (‘Sonderforschungsbereich’; SFB) 665 and 958 and Charité Promotionsabschlussstipendium.

## Author Contributions Statement

SAS, SAK, and HZ designed the study. HZ carried out the experiments. BS prepared primary neurons; BS, NR, and SAK assisted with data analysis. HZ and SAS wrote the manuscript text and HZ prepared figures. All authors reviewed the manuscript.

## Additional Information

The authors declare no competing interests

