## Supplement for "Disease-associated scaffold protein CNK2 modulates PSD size and influences trafficking of new synaptic binding partner TNIK"

Charité-Universitätsmedizin Berlin

Charitéplatz 1

10117 Berlin

GERMANY

### Supplement:

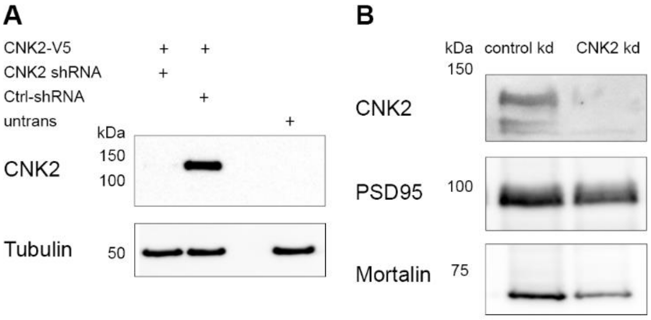


**Supplement Figure 1: shRNA mediated knockdown of CNK2**

(A) Western blot of shRNA mediated knockdown of V5 tagged CNK2 expressed in COS-7 cells. Western blot was probed with anti-V5 antibody. Tubulin serves as loading control. (B) Synaptosome preparation of primary neurons (DIV 23) infected with lentivirus transducing shRNA (DIV 3) to knockdown endogenous CNK2. Control knockdown (left lane) CNK2 knockdown (right lane) tested by western blot with antibody detecting endogenous CNK2 (upper panel). PSD-95 (middle panel) and Mortalin (lower panel) serve as loading control.

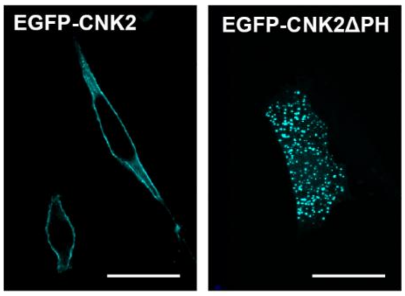


**Supplement Figure 2: The CNK2 variant CNK2ΔPH does not bind to the membrane**

Image of COS-7 cells overexpressing EGFP-CNK2 (left) or EGFP- CNK2ΔPH (right) Scale bar: 20µm

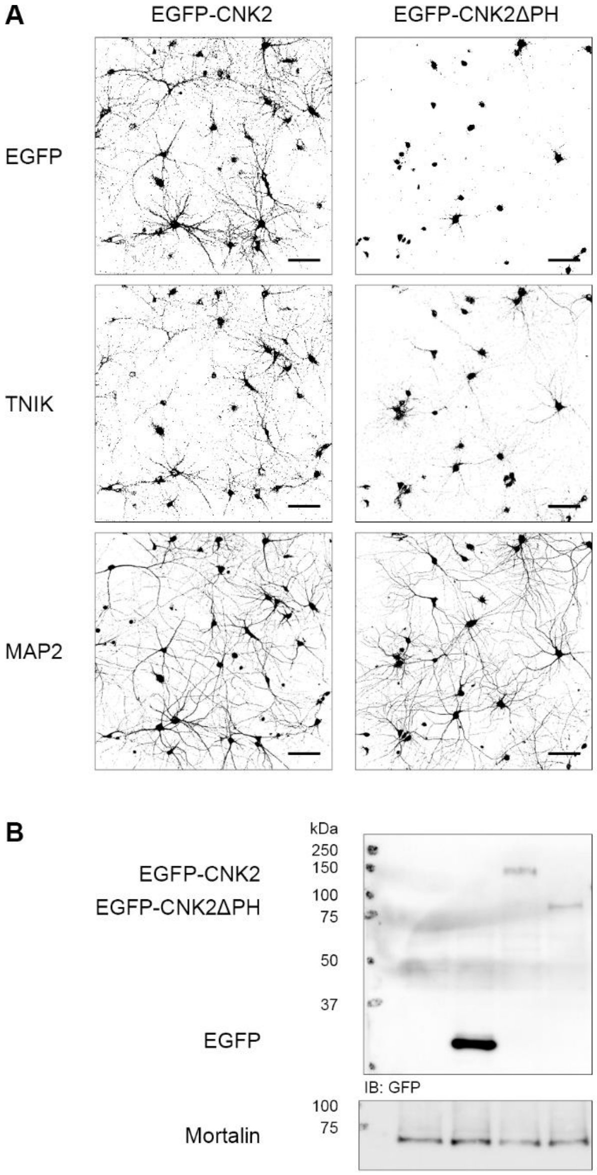


**Supplement Figure 3: CNK2ΔPH is mis-localised in neurons**

(A) Overview images of neurons (DIV 23) expressing EGFP-CNK2 or EGFP-CNK2ΔPH following viral-mediated gene delivery. Lower panel depicts MAP2, indicating the general dendritic structure of neurons. Thresholds are set identical for each panel (EGFP/TNIK/MAP2). Scale bar: 100 µm. (B) Western blot of whole cell lysates from cultured hippocampal neurons infected with EGFP, EGFP-CNK2 or EGFP-CNK2ΔPH; proteins detected by western blot with anti-GFP antibody. Mortalin as loading control.

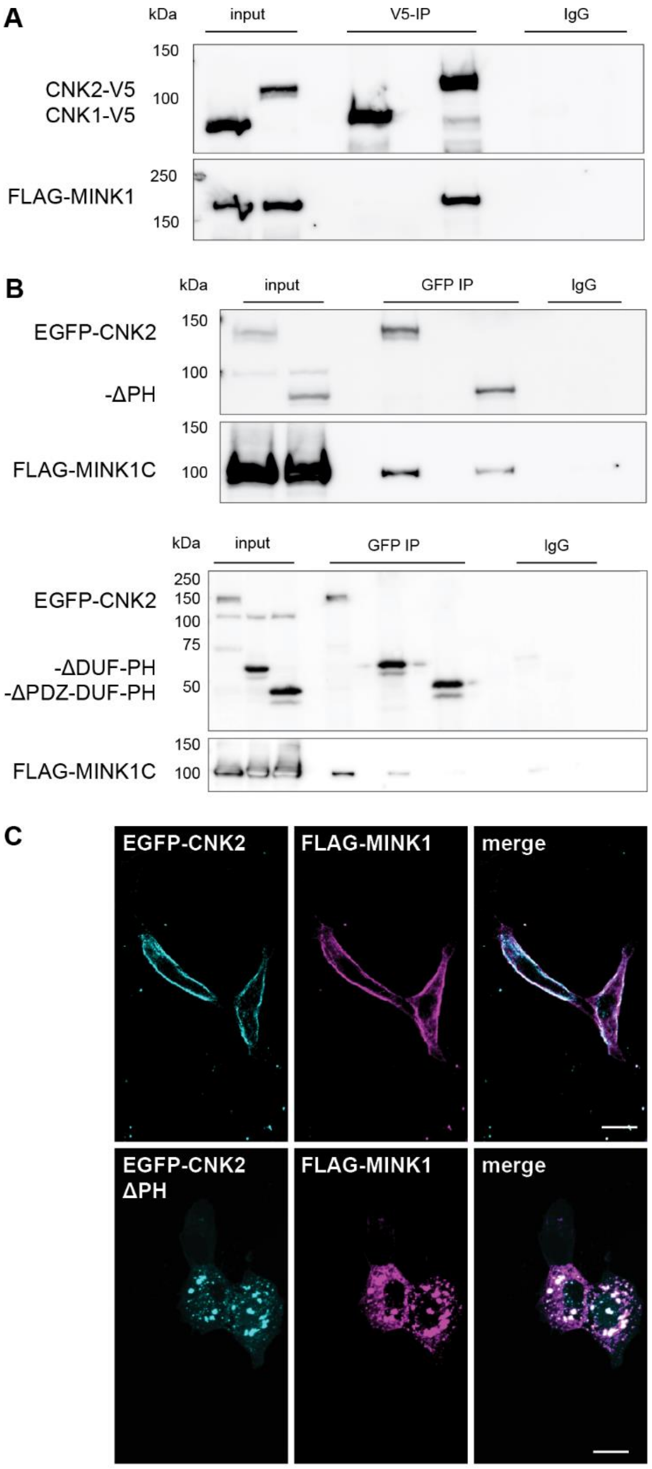


**Supplement Figure 4: CNK2 interacts with MINK1 and regulates its localisation**

(A) CNK2 specifically interacts with MINK1. Co-immunoprecipitation experiment with CNK1-V5, CNK2-V5 and FLAG-MINK1 overexpressed in COS-7 cells. Proteins were immunoprecipitated with either anti-V5 (mouse) antibody or normal mouse IgG as a negative control. (B) Co-immunoprecipitation experiments of EGFP-CNK2 variants overexpressed in COS-7 cells together with FLAG-MINK1C (aa 534-1301). Proteins were immunoprecipitated with either anti-GFP (mouse) antibody or normal mouse IgG as negative control. Proteins were detected by WB. Input control (lysate) is on the left. (C) Immunofluorescence experiment in COS-7 cells expressing FLAG-MINK1 together with EGFP-CNK2 (upper panel) or EGFP-CNK2ΔPH (lower panel); Left lane shows EGFP-tagged CNK2 variants (cyan), middle lane shows FLAG-MINK1 (magenta), and right lane shows merged channels. Scale bar: 10 µm

| Gene Name | Gene ID | Interaction domain (AA) | Clones detected |
|---|---|---|---|
| Arhgap39 | 223666 | 1-319 | 2 |
| Cytohesin1 | 19157 | 14-100 | 7 |
| Cytohesin4 | 72318 | 6-235 | 5 |
| Magi3 | 99470 | 63-656 | 1 |
| Mink1 | 50932 | 430-533 | 4 |
| Rlf | 109263 | 989-1496 | 1 |
| Samd12 | 320679 | 41-161 | 3 |
| Sox5 | 20678 | 351-582 | 1 |
| Tnik | 665113 | 337-598 | 1 |

**Supplement Table 1: Yeast-two-hybrid screen of cDNAs form adult mouse brain**

The bait full-length CNKSR2 (mus musculus, Gene ID: 245684, aa 1-1032), cloned into pB27 (N-LexA-bait-C fusion) was used with an adult mouse brain cDNA library

**Supplement Fig. 5: full blots**

**Figure 1A**

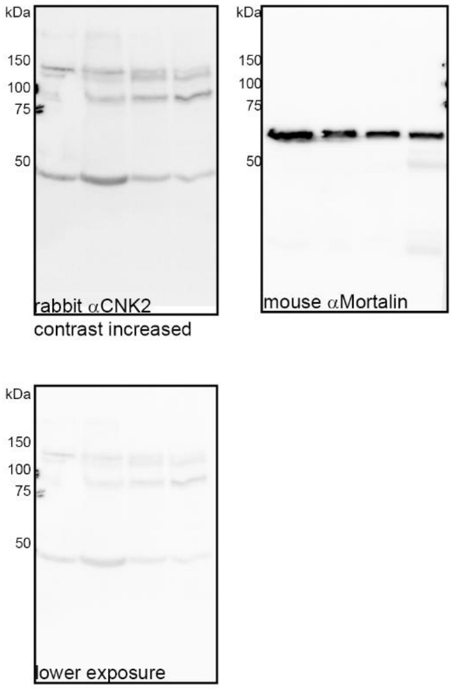


**Figure 4B**

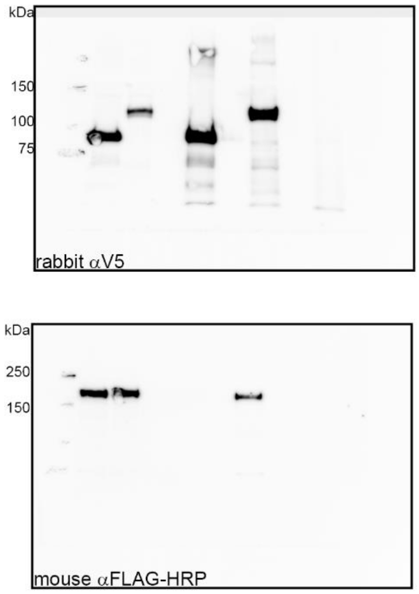


**Figure 4D**

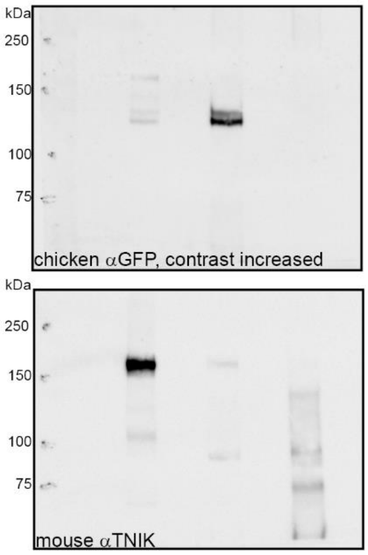


**Figure 5B**

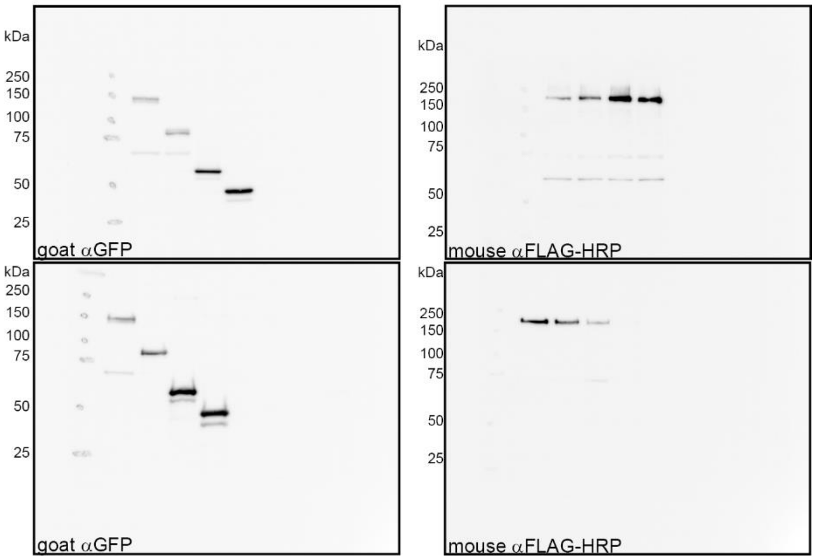

**Supplement Fig. 5: full blots**

**Supplement Figure 1A**

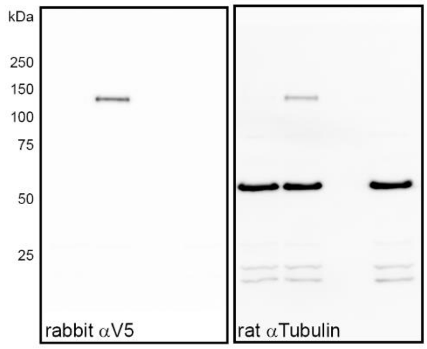


**Supplement Figure 1B**

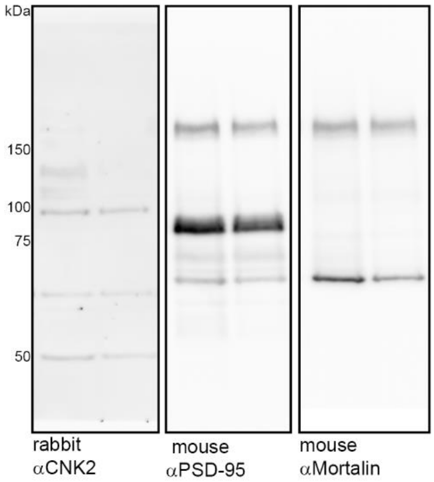


**Supplement Figure 3B**

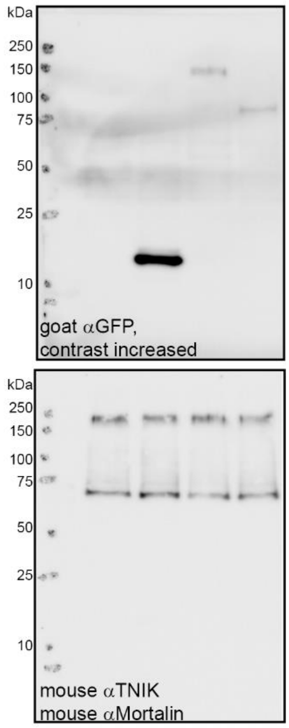


**Supplement Figure 4A**

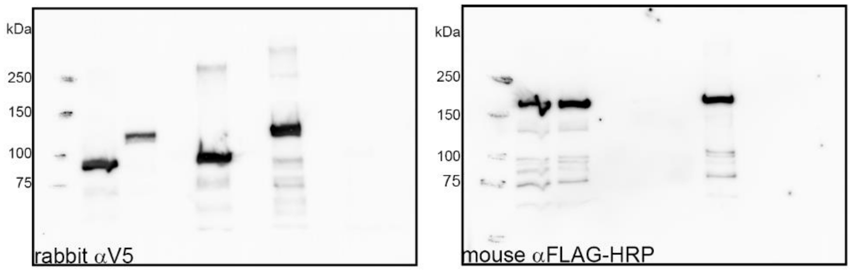


**Supplement Figure 4B**

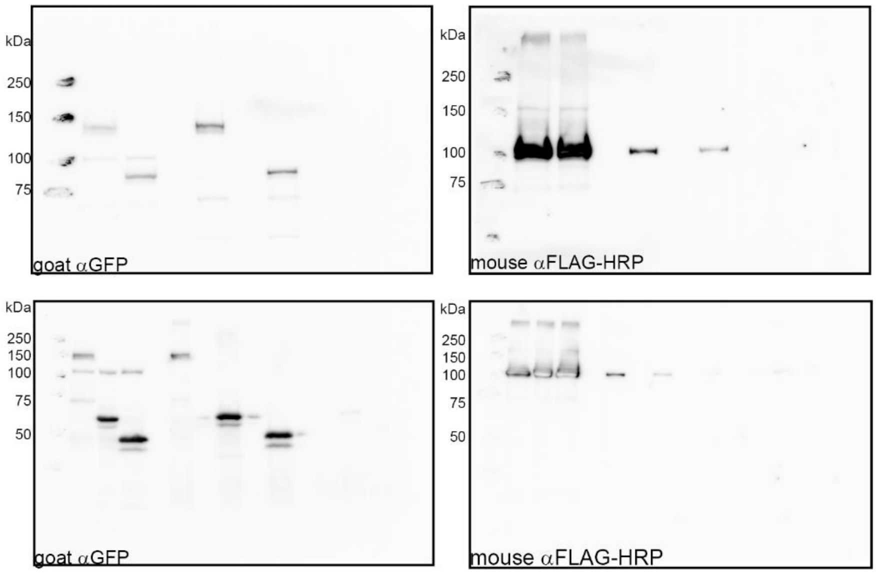
